# Analysis of isoform complexity in pan-transcriptome graphs with atroplex

**DOI:** 10.64898/2026.06.30.735696

**Authors:** Richard A. Schäfer, Yangyang Li, Joshua Fry, Rendong Yang

## Abstract

**Motivation:** Alternative splicing of precursor mRNA lets a single gene encode multiple isoforms by joining exons in different combinations. Long-read sequencing resolves this isoform diversity across tissues, cohorts, and conditions. However, the resulting pan-transcriptomes are structurally complex, and their analysis requires repeatedly searching the full catalogue, which is impractical without a queryable index. As splicing patterns differ across conditions, a structure is needed that captures the connectivity between exons, not just their coordinates, so isoforms can be compared by structure across cohorts.

**Results:** We present `atroplex`, a framework that indexes pan-transcriptome annotations and transcript isoforms in a combined spatial index and graph overlay, capturing both exon coordinates and splice connectivity. `atroplex` classifies query transcripts against the index, tracks per-sample isoform presence, and enables cross-cohort isoform comparison. We indexed 21,005 samples spanning multiple reference resources into a single queryable structure, yielding a comprehensive map of isoform complexity that supports improved transcript discovery and structural comparison across cohorts.

## Introduction

Alternative splicing (AS) is an eukaryotic cellular process in which multiple mature mRNAs can be produced from the same pre-mRNA by the different inclusion or exclusion of exon segments. It is estimated that roughly 95% of multi-exon genes undergo alternative splicing, affecting the vast majority of protein-coding genes (Pan et al., 2008). Through this mechanism, a single gene can give rise to protein isoforms with distinct domains, cellular localizations, and interaction partners, thereby expanding proteomic diversity without increasing genome size. The dysregulation of AS has been implicated in numerous diseases, especially neurodegenerative disorders (Nikom and Zheng, 2023), autoimmune conditions (Ren et al., 2021) and cancers (Bradley and Anczuków, 2023). Consequently, there is a critical need to study AS at isoform resolution and across large cohorts to understand how specific transcript structures contribute to physiology and disease. In that regard, long-read sequencing enables the direct observation of full-length isoforms and their structural diversity (Amarasinghe et al., 2020). However, the analysis of transcript architectures across multiple datasets remains challenging. Available tools such as `Bambu` (Chen et al., 2023), `FLAIR` (Tang et al., 2020), `TALON` (Wyman et al., 2020), and `IsoQuant` (Prjibelski et al., 2023) focus on transcript discovery and quantification within a single dataset. `SQANTI3` (Pardo-Palacios et al., 2024) categorizes transcripts against a reference but operates per-sample without population context. The LRGASP consortium revealed substantial variability between these methods, both in the number and identity of predicted transcript models, with limited isoform overlap and low quantification concordance (Pardo-Palacios et al., 2024). More recently, `isopedia` (Zheng et al., 2026) demonstrated the value of population-scale isoform indexing by cataloging read-level splice junction signatures. This enables recurrence-based distinction of known and novel isoforms. However, its read-level representation lacks the exon connectivity needed to analyze how transcript structures diverge at specific splice sites to characterize splicing complexity across cohorts. Thus, no existing framework combines structural graph representation with population-scale indexing to enable splicing complexity analysis across cohorts. The underlying problem of the query and storage of genomic intervals is crucial across genomics applications. For example, in homology detection to find conserved sequences across species (Lott et al., 2018; Schäfer et al., 2020), in RNA-RNA interaction prediction to pinpoint binding sites within specific gene regions, such as UTRs (Schäfer and Voss, 2021), or retrieval of overlapping somatic variants to assemble candidate peptides from RNA-seq data (Schäfer et al., 2023). Transcript isoforms are collections of exon intervals linked by splice junctions, and querying them at scale requires efficient spatial indexing combined with graph traversal. Recently, we evaluated existing tools for querying genomic intervals (Schäfer and Yang, 2025) and showed that implicit-interval-tree-based methods such as `bedtk` (Li and Rong, 2020) are efficient for simple overlap queries, but are not able to perform complex chained lookups. In contrast, tree-based indexing structures such as `GIGGLE` (Layer et al., 2018) support complex queries but are inefficient for heterogeneous intervals produced by long-read RNA-seq, and none expose a typed graph overlay that would allow transcripts, segments, and exons to be traversed as linked structures rather than unrelated intervals. To address this gap we developed `atroplex`, a pan-transcriptome analysis toolkit built on `genogrove` (https://github.com/genogrove), a high-performance genomic indexing library that combines an interval B+ tree with a typed graph overlay. This allows (i) the construction of a unified pan-transcriptome index from heterogeneous annotations, analysis of structural sharing, splicing complexity, and per-sample diversity, (iii) classification of transcripts against the index, and (iv) discovery of novel transcripts from aligned long reads.

## Results

### Pan-transcriptome indexing with **atroplex**

We built `atroplex` on `genogrove`, that is a general-purpose genomics indexing library we developed, pairing a graph layout with an integrated spatial index (Figure 1). Keys in a grove are genomic intervals carrying arbitrary feature data that are connected by directed edges. A subset of these keys is organized in an interval B+ tree with configurable order, while keys reached only through edge traversal are registered as external keys outside the tree. Spatial range queries return overlapping keys as entry points into the graph, from which filtered edge traversal walks a specific path. These groves can be built from sorted intervals or by single-key insertion (Supplementary Figure S1). In `atroplex`, we store the transcript structure, that is the ordered chain on a given strand and chromosome as a single spatially indexed segment, and its exon chain is stored as external keys linked through directed edges. This separation enables a two-phase query strategy: a coarse spatial intersection retrieves candidate segments overlapping a genomic region, followed by fine-grained graph traversal along exon chains for structural comparison. Because each edge carries the segment’s numeric index as its identifier, paths can be unambiguously resolved at branching exons where multiple segments share an exon node but diverge downstream. Transcripts from multiple samples and annotation sources are integrated into a single index, one for each chromosome. The segments that share identical exon structures across files are merged into a single entry, with sample provenance on each feature. Transcripts whose exon chain is a contiguous ordered subsequence of an existing segment are absorbed into the longer parent, rather than creating a separate entry (Supplementary Figure S2). Absorption candidates are identified spatially via the grove’s interval index, making the process independent of gene identifier conventions across input files and enabling seamless integration of heterogeneous annotation sources. Per-sample expression values are stored separately, enabling the index to scale to tens of thousands of samples. We applied this construction to a comprehensive short and long-read pan-transcriptome built from 21,005 samples spanning multiple reference resources: The GENCODE reference annotation (Harrow et al., 2012), ENCODE long-read RNA-seq (*n* = 144 samples across 55 biosample groups) (Reese et al., 2023), TCGA cancer cohorts (*n* = 11,096 across 33 cancer types), and GTEx tissue cohorts (*n* = 9,765 across 31 tissues). `StringTie2` (Kovaka et al., 2019) was used to assemble both the TCGA and GTEx cohorts.

**Figure 1.**
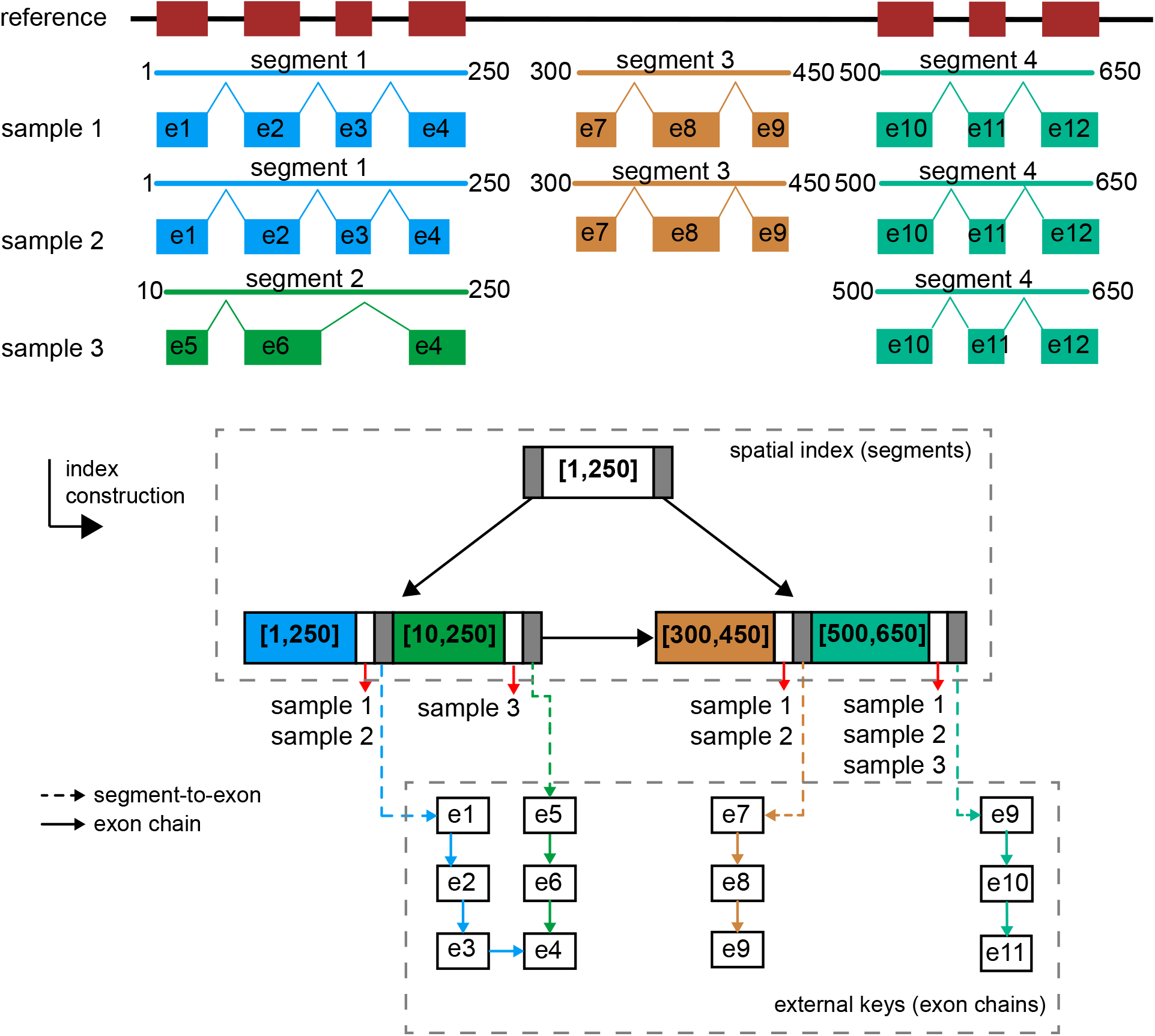
Overview of the atroplex workflow. `atroplex` layers a pan-transcriptome construction and analysis workflow on top of genogrove: transcripts from heterogeneous annotation and assembly inputs are decomposed into exon chains, deduplicated by exact-match and absorbed by ISM rules, and indexed as segments (spatial keys) connected to their constituent exons (graph nodes), yielding a single grove that supports per-sample provenance, structural classification, and splicing-graph traversal in one pass.

The resulting index contains 3,468,212 deduplicated segments built from 2,674,603 unique exons across 25 chromosomes, with 269,785,338 graph edges encoding splice connectivity (Figure 2A). Per-source contribution analysis (Figure 2B) shows that HAVANA contributes 562,588 segments (62.6 % source-exclusive) and ENSEMBL 22,453 (50.9 % exclusive), together forming the curated annotation backbone. `StringTie2` contributes 3,076,747 segments (92.8 % source exclusive) reflecting the dominance of cohort-derived novel structures at cohort scale. TALON that was used to process the ENCODE datasets contributes 31,166 segments (85.0 % source-exclusive), recapitulating the long-read pipeline’s tendency to discover novel splice combinations among already-annotated exons. Of the 3.47 million indexed segments, 2,233 are present in ≥95 % of all 21,005 samples and define a soft-core pan-transcriptome (Figure 2C). This conservation core is dominated by protein-coding biotypes (1,875*/*2,233 = 91.3 %), with smaller contributions from processed pseudogenes (5.5 %) and Mt_tRNA (1.0 %). Notably, 179 conserved segments (8.0 %) are located on novel-locus genes carrying no GENCODE biotype annotation.

**Figure 2.**
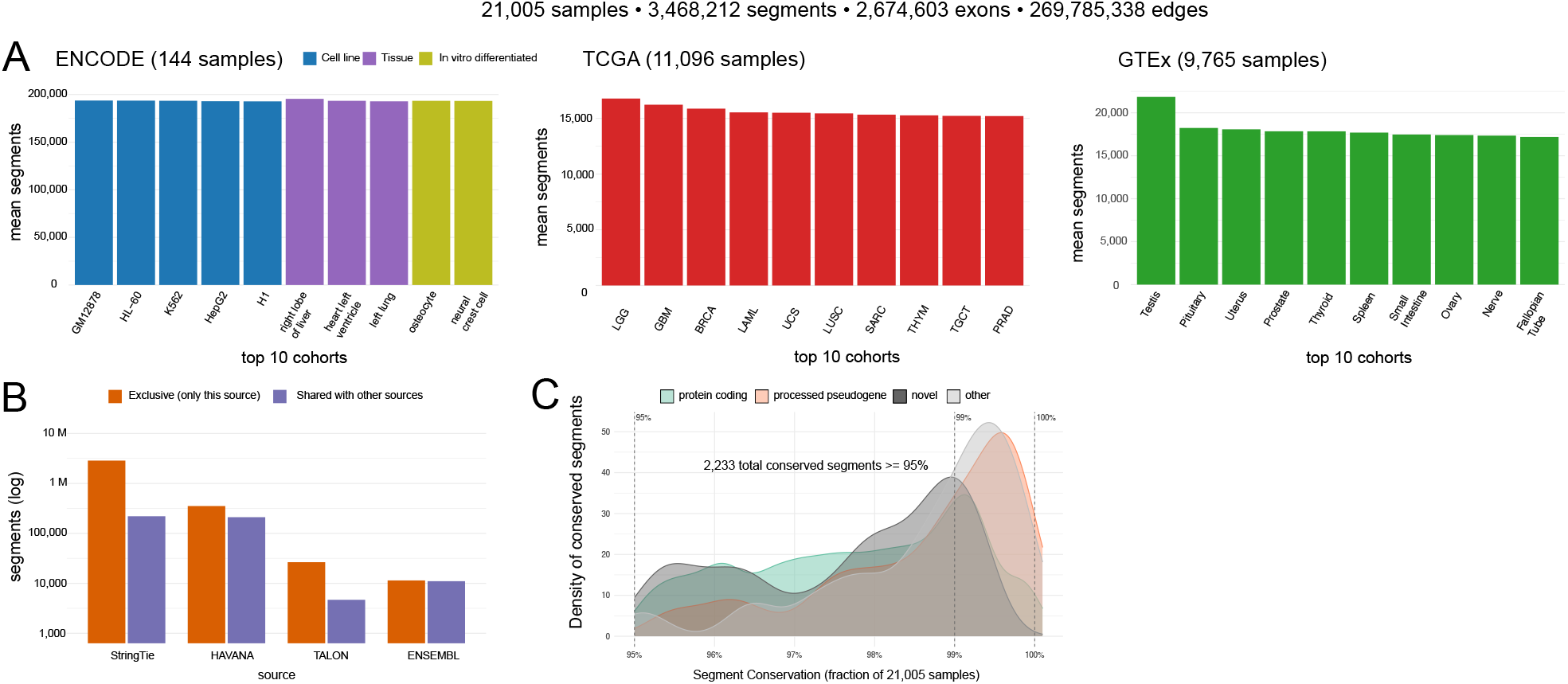
Composition of the atroplex index. The index aggregates 21,005 samples spanning three reference resources that result in 3,468,212 segments, 2,674,603 exons, and 269,785,338 graph edges **(A)**For each dataset, the ten cohorts contributing the most segments on average are shown; `ENCODE` (*n* = 144 samples across 55 biosample groups: cell lines, tissues, primary cells, and `in vitro` differentiated cells); `TCGA` (*n* = 11,096 samples across 33 cancer cohorts); `GTEx` (*n* = 9,765 samples across 31 tissues). **(B)**Per-source contribution to the indexed segments. The 3,468,212 indexed segments were grouped by the annotation/assembly source (GFF/GTF column 2). These include manual (HAVANA) and automated annotation (ENSEMBL), short-read assembly (StringTie), and long-read isoform calling (TALON). For each source, segments are partitioned into source-exclusive (present only in that source) and shared (also present in *≥* 1 other source). (**C**) Segment conservation distribution. 2,233 segments are present in ≥ 95% of the 21,005 samples. For each segment, the fraction of samples carrying the segment (conservation depth), broken down by biotype is shown.

**Figure 3.**
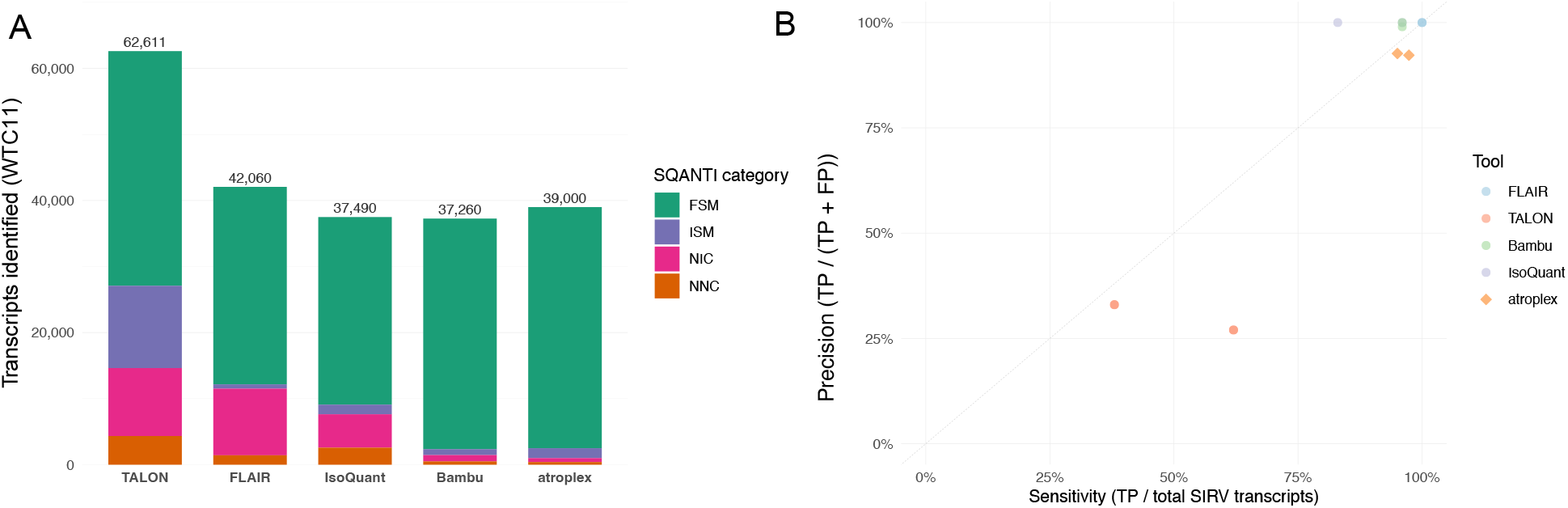
Benchmarking isoform discovery. **(A)** Structural-category breakdown of isoform discovery on the WTC11 PacBio long-read benchmark. Stacked bar chart showing the median number of transcripts identified per tool across LRGASP PacBio pipelines. **(B)** Sensitivity versus precision on SIRV synthetic spike-in transcripts.

### Structural analysis of splicing complexity across samples

To characterize the structural landscape of the pan-transcriptome, `atroplex` performs a single-pass traversal of the indexed grove (`atroplex` inspect). This reports an overview of the index composition such the number of segments and their distribution per sample, source and biotype. In addition, sharing statistics explore to which extend the segments and its exons are shared between samples or are exclusive. This also creates a splicing event catalog that classifies the exons in the samples as either cassette exons, alternative 5^*′*^/3^*′*^ splice sites, intron retention, mutually exclusive exons, and alternative terminal exons with per-sample Percent Spliced in (PSI) (Schafer et al., 2015). Finally, the splicing hub analysis identifies exons with substantial distinct downstream targets as decision points for complex alternative splicing. This is quantified by per-sample PSI and Shannon entropy of the branch usage distribution (Sterne-Weiler et al., 2018). This was applied using default parameters to our indexed pan-transcriptome (Figure 4).

**Figure 4.**
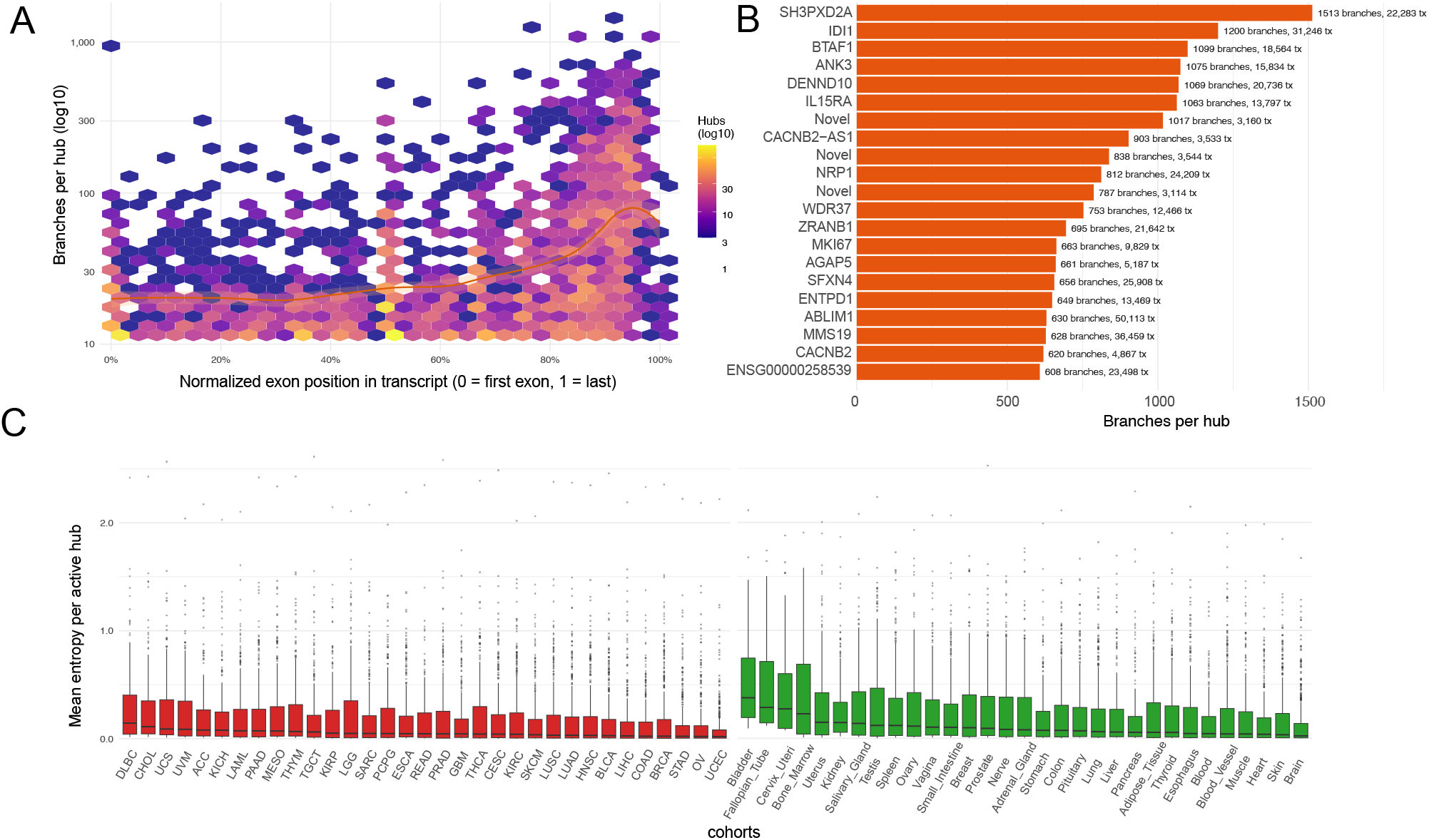
Isoform complexity across cohorts. (**A**) Splicing-hub branching capacity by normalized exon position. Density of exons with multiple distinct downstream branch targets (splicing hubs) plotted against the normalized exon position. (**B**) Top splicing hubs by branching capacity, labeled by parent gene name and annotated with total branch count and the number of source transcripts. (**C**) Per-cohort splicing-hub entropy distribution. Boxplots of cohort-median entropy across active splicing hubs (entropy *>* 0) for each of 64 short-read StringTie cohorts (33 TCGA cancer cohorts; 31 GTEx tissue cohorts), faceted by dataset.

### Transcript classification reveals differential isoform usage across sample groups

To assess how individual transcripts relate to the indexed pan-transcriptome, `atroplex` is able to classify query transcripts against the grove (`atroplex` query), following the classification introduced in `SQANTI` (Pardo-Palacios et al., 2024). These include *Full Splice Match (FSM), Incomplete Splice Match (ISM), Novel in Catalog (NIC), Novel not in Catalog (NNC)*, as well as *genic intron* (mono-exon within intron), *genic-genomic* (overlaps both intron and exon), *antisense* (opposite strand overlap), and *intergenic* (no overlap). At first, a spatial query on the interval tree retrieves the candidate segments that overlap the coordinate span of the query transcript which is followed by graph traversal along the candidates exon chain for structural comparison. In the following, the junctions in the query transcripts are compared against the candidate’s exon boundaries, and the best matching segment determines the structural category. For each classified transcript, the output reports per-sample presence across the cohort and, when available, per-sample expression at the matched locus. Because classification against the pan-transcriptome yields per-sample expression for every matched segment, differential transcript usage (DTU) can be tested directly without re-quantifying reads from raw alignments. Given a two-group contrast defined by the manifest’s group annotation (`--contrast`), the mean expression of each segment is computed per group, converted to within-gene proportions, and tested for differences using a gene-level *χ*^2^ statistic on the 2 ×*k* contingency table (two groups by *k* segments per gene), with *p*-values corrected by the Benjamini– Hochberg procedure. This identifies genes whose isoform repertoire shifts between conditions and reports which specific isoforms drive the differential usage.

### Structural transcript discovery improves read-level isoform resolution

In transcript discovery from long-reads, individual read alignments are typically compared against a flat reference annotation, classifying each as matching or novel without capturing how a novel isoform relates to known splicing paths. As the pan-transcriptome captures the full splicing topology across cohorts, it provides a richer structural reference: novel reads can be characterized not only as structurally distinct but by where they diverge from the population’s known splice decisions. To this end, `atroplex` extracts splice junctions from aligned reads clustered by compatible junction signatures (`atroplex` discover). Each cluster enters the grove through a spatial query and is walked along the exon graph for structural comparison. Clusters that follow an existing path are recorded as read support for that isoform, while clusters that diverge are characterized by their point of divergence — at a splicing hub, a novel exon boundary, or a skipped cassette — and inserted as new segments. This enables iterative index refinement: transcripts discovered from one cohort become part of the reference for subsequent analyses.

### Isoform complexity across cohorts

The splicing-hub analysis identified 2,577 splicing-decision exons in 761 named genes across the index. Splicing capacity (number of distinct downstream branches per hub) ranges from 3 to 1,513 alternative targets and is strongly biased toward the 3^*′*^ end of transcripts (Figure 4A): median branching capacity rises monotonically from 16 alternative targets in the first decile of normalised exon position to 80 in the last decile — a five-fold enrichment recovering the well-established biology of alternative last-exon usage and alternative polyadenylation. The most splicing-rich genes (Figure 4B) include canonical alternatively-spliced loci, each carrying ≥ 11 distinct splicing hubs. At the cohort level (Figure 4C), per-cohort splicing-hub entropy distributions across the 64 short-read StringTie cohorts (33 TCGA cancer cohorts and 31 GTEx tissues) reveal modest but consistent shifts in active-hub branch diversity, capturing tissue- and cancer-specific splicing-decision patterns that emerge only at cohort scale.

## Discussion

`atroplex` addresses a gap between per-sample transcript discovery tools and population-scale isoform catalogues by representing the pan-transcriptome as a combined spatial index and typed graph overlay. By storing transcript structure as spatially indexed segments linked to their exons through directed edges, the framework supports both coarse coordinate queries and fine-grained structural comparison in a single pass, while keeping per-sample provenance and expression at cohort scale. Indexing 21,005 samples into one queryable structure revealed a soft-core pan-transcriptome dominated by protein-coding biotypes and a 3^*′*^-biased landscape of splicing-decision hubs that recovers known alternative last-exon and polyadenylation biology. Because classification yields per-sample expression for every matched segment, differential transcript usage can be tested directly from the index without re-quantifying raw reads, and discovery clusters that diverge from known paths can be reinserted to iteratively refine the reference. We anticipate that this structural, population-aware view of isoform complexity will support transcript discovery and cross-cohort comparison in settings where flat per-sample annotations fall short.

## Conflicts of interest

The authors declare that they have no competing interests.

## Funding

This project is supported in part by NIH grants R35GM142441 and R01CA259388 awarded to R.Y.

## Data availability

The ENCODE4 long-read RNA-seq samples used in this study are publicly available from the ENCODE Portal (https://www.encodeproject.org). All experiment accessions, biosample metadata, replicate structure, and expression units for the samples indexed with `atroplex` are listed in Supplementary Table 1. Reference annotations are from GENCODE v49 on the GRCh38 assembly.

## Author contributions statement

R.A.S. and Y.L. designed and implemented the software. R.A.S. performed the benchmarks. J.F. curated and processed the data. R.Y. acquired funding and supervised the project. R.A.S. and R.Y. wrote the manuscript. All authors read and approved the final manuscript.

## Acknowledgments

The results shown here are in whole or part based upon data generated by the TCGA Research Network: https://www.cancer.gov/tcga and the Genotype-Tissue-Expression (GTEx) Portal (https://gtexportal.org). We thank Fikrat Talibli for valuable discussions on the graph-based routines underlying `genogrove`. During the preparation of this manuscript, LLMs were used for code review and language editing.

## Supplementary Information

### The genogrove data structure

`genogrove` (https://github.com/genogrove) provides a parameterized container that fuses two indices into one addressable object: an interval B+ tree of configurable order *k* keyed on a genomic coordinate type, and a typed graph overlay whose nodes are the keys held by the tree. Spatial lookup over *n* keys runs in *O*(log_*k*_ *n* + *m*) for *m* reported intervals; graph traversal from any node visits each incident edge once. Both indices share their key storage, so a spatial query can hand its hits directly to a graph walk without copying or re-resolving keys, and a single in-memory image is sufficient for both forms of access.

For the pan-transcriptome analysis, the grove is instantiated with strand-aware genomic coordinates as keys and a feature type that distinguishes segments from exons. Segments are inserted as ordinary keys and therefore participate in spatial queries, while exons are registered as external keys, reachable only by traversing an edge from a segment and never returned as spurious hits on a coordinate intersection. Each edge carries a numeric segment identifier together with an edge-type label (SEGMENT-TO-EXON, EXON-TO-EXON, or SEGMENT-TO-SEGMENT). Once candidate segments have been identified in the spatial query, the SEGMENT-TO-EXON edge to the first exon is followed, then the EXON-TO-EXON edges are walked in 5^*′*^-to-3^*′*^ order along the chain. Edges are deduplicated by reusing each segment’s identifier as the edge label: the first transcript to materialize a given exon chain creates the segment and its full chain of edges, and any later transcript with the identical chain reuses the segment and skips edge creation entirely, so edge construction scales with the number of *distinct* splice chains rather than with the number of input transcripts. When two structurally distinct segments share an exon pair *A* → *B*, each inserts its own *A* → *B* edge tagged with its own segment identifier; traversal from any starting segment then disambiguates the correct downstream path at a branching exon by filtering on the current segment’s identifier. This is what allows the overlay to behave as a multigraph rather than as a simple graph, and it is the property that keeps exon-chain reconstruction a deterministic linear walk even at high-fan-out hubs.

To keep the index memory efficient at cohort scale, every repeated string in the input (e.g., gene and transcript identifiers, annotation source labels, sample names) is interned through registries and replaced on the feature by an integer handle. The annotation provenance (e.g., HAVANA, ENSEMBL, TALON) is encoded as a small bitfield. Similarly, the sample membership uses a bitset to support thousands of samples. The transcript ids are stored as compact vectors and the expression values are streamed to quantification sidecars at build time, and read back on demand (subcalls `inspect and query`).

### Construction of the pan-transcriptome index

The build proceeds gene-by-gene over the input annotation and sample files. Reference annotations are processed before experimental samples so that annotation segments are present in the index before sample transcripts are matched against them. Transcripts are sorted by exon count in descending order so that longer isoforms are inserted first and remain available to absorb shorter ones. For each transcript, two types of features are contributed to the grove: the full coordinate span is registered as a spatially indexed segment, and the individual exons are added as graph-only nodes that participate in traversal but not in coordinate queries. Exons are deduplicated against a per-chromosome coordinate cache so that boundaries shared across files or transcripts collapse to a single node. The segment is then linked to its first exon by a SEGMENT-TO-EXON edge, and successive EXON-TO-EXON edges chain the exons in 5^*′*^-to-3^*′*^ order; each edge carries the originating segment’s identifier so that traversal at branching exons remains unambiguous.

Before a new segment is created, the transcript’s exon chain is evaluated against the existing index in two stages. An exact structural match is first probed via a hash of the ordered exon coordinates, in which case the matching segment’s metadata is updated in place. If no exact match is found, the spatial index is queried for segments whose coordinate span overlaps the transcript on the same strand, and the candidates are evaluated against a set of absorption rules that decide whether the incoming transcript should be merged into an existing segment or retained as a distinct isoform (Supplementary Figure S2). Transcripts whose exon chain is a contiguous ordered subsequence of an existing segment are classified by their truncation pattern: 5^*′*^ ISMs, intact at the 3^*′*^ end but missing one or more exons at the 5^*′*^ end, are retained as separate segments because they may reflect genuine alternative transcription start sites; 3^*′*^ISMs, missing one or two 5^*′*^ exons, are absorbed into the parent because they typically arise from long-read truncation artefacts; and internal fragments, truncated at both ends, are dropped when the parent derives from a reference annotation but retained when it derives from a sample, on the rationale that fragments matching reference internals are unlikely to be biologically informative whereas fragments co-occurring across samples may signal genuine partial assemblies. Terminal variants — transcripts sharing an identical intron chain with a catalogued segment but differing by fewer than 50 bp at the TSS or TES — are likewise absorbed. When exact coordinate comparison fails, a fuzzy pass tolerates up to a configurable boundary difference (5 bp by default) to accommodate minor discrepancies between annotation versions or assembly pipelines.

When a transcript passes all absorption checks without matching an existing segment, a new segment is created and inserted into the spatial index. If the new segment spatially overlaps an annotation segment, it inherits the annotation’s gene identity, preventing sample-specific identifiers from inflating the gene count. A reverse absorption pass then checks whether any previously inserted segments are contiguous subsequences of the new one and merges them into the longer parent. Mono-exon transcripts from experimental samples are classified by spatial overlap with the existing index: those spanning from one exon across an intron into the next exon of a multi-exon segment are retained as candidate intron-retention events, while those overlapping only exonic regions or falling in intergenic space are discarded. Mono-exon transcripts from reference annotations are always retained to preserve the completeness of the reference gene catalog.

We applied this procedure to construct a pan-transcriptome index from 21,005 samples spanning the ENCODE4 long-read RNA-seq collection (Reese et al., 2023), the GTEx tissue atlas (Lonsdale et al., 2013), and the TCGA cancer cohort (Weinstein et al., 2013) (Supplementary Figure S4). For the GTEx and TCGA cohorts, transcript assemblies were generated from aligned BAM files with StringTie v2 (Kovaka et al., 2019) in conservative mode. For the ENCODE4 long-read samples, we used the TALON-processed GTF files provided by the consortium and injected per-transcript read counts from the corresponding TALON quantification tables. The build used GENCODE v49 as the reference annotation, restricted the index to isoform variation within annotated gene loci, applied expression thresholds of ≥10 counts, ≥10 coverage, and ≥5 TPM to exclude low-confidence transcripts, and required each isoform to be present in at least half of the biological replicates within its experimental group.

### Segment absorption

Long-read sequencing routinely produces incomplete splice matches that are transcripts whose exon chain is a contiguous ordered subsequence of a longer transcript, typically arising from 5^*′*^ or 3^*′*^ truncation during library preparation or base-calling. If such transcripts are admitted to the index as independent segments, they fragment what should be a single isoform into a family of nested variants. `atroplex` therefore attempts, at segment creation time, to absorb each incoming transcript into an existing segment of the same gene whose exon chain contains the new chain as a contiguous subsequence.

Subsequence checking is performed in a per-gene, per-chromosome index that stores for every live segment its ordered vector of exon-chain keys. Candidate parents are enumerated in gene order, and containment is tested in two passes: an exact-coordinate pass first, followed by a fuzzy pass that tolerates up to `--fuzzy-tolerance` bp of mismatch (default 5 bp) on the first and last internal boundaries to accommodate minor assembly differences between reference and sample annotations. If an incoming transcript matches a parent, its transcript ID, sample bits, source bits, and per-sample expression are merged into the parent rather than creating a new segment. In the case of a longer transcript arriving after a shorter one it would be given preference. In the reverse absorption, the shorter segment is merged into the longer one, and the shorter segment is marked as *absorbed* via a boolean tombstone flag. All downstream consumers (statistics, classification, discovery, export) filter out features with the tombstone set, and the tree entries themselves will be removed. Mono-exon transcripts are never considered either as absorption candidates or as parents, because with only a single exon boundary the parent assignment is ambiguous. Absorption can be disabled globally with `--no-absorb`, in which case each transcript produces its own segment. Supplementary Figure S2 illustrates the absorption rules, and Supplementary Figure S3 the decision flow during segment creation.

### Streaming pan-transcriptome statistics

`atroplex` computes statistics at two tiers. A lightweight summary is reported during the grove construction directly from the builder cache, regardless of the subcommand invoked; it contains overview counts, biotype breakdowns, per-chromosome distributions, and sample counts. A *full analysis report*, produced by the `inspect` subcommand, performs a single-pass streaming traversal of the grove and writes a tree of text, CSV, and TSV files under category subfolders `(overview/, sharing/, splicing_hubs/, splicing_events/)`. Counters are maintained per sample as flat vectors rather than as nested maps, which keeps peak memory linear in the number of samples and avoids the quadratic blow-up of earlier per-pair diversity metrics on cohorts with more than 10^3^ samples.

Sharing statistics are defined as follows. An exon or segment is *exclusive* if exactly one sample bit is set in its `sample_bitset`; *shared* if two or more bits are set; and *conserved* if all sample-type entries in the manifest carry a bit. At the exon level we additionally distinguish *constitutive* exons (used by every transcript of their parent gene) from *alternative* exons. Per-gene isoform diversity is summarized by the Shannon entropy of the exon-usage distribution,

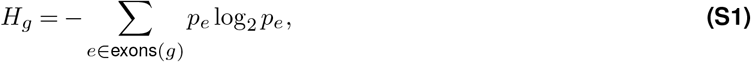

where *p*_*e*_ is the fraction of transcripts of gene *g* that use exon *e*, normalized so that *p*_*e*_ = 1. The effective isoform count 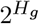 reports how many equally used isoforms gene *g* would need to explain the observed exon usage if they were uniform; it is scale-invariant and comparable across genes of different sizes. Both metrics are *O*(*N*_*g*_) per gene and require no pairwise transcript comparison.

### Splicing hubs

We define a *splicing hub* as an exon with strictly more than MinHubBranches = 10 distinct downstream exon targets across all samples in the index. For each hub exon in each sample, we compute the Percent Spliced In (PSI) of each downstream branch as

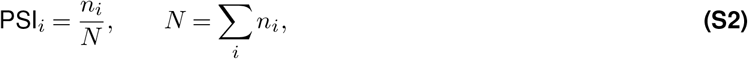

where *n*_*i*_ is the number of transcripts that traverse the hub exon and continue to downstream target *i*. The splicing entropy at the hub in that sample is then

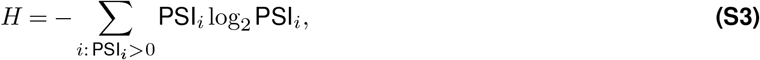

which is zero if all transcripts at the hub take a single path and reaches its theoretical maximum log_2_ *k* (where *k* is the number of downstream targets) when branch usage is uniform. Per-hub output records, in addition to *H*, the number of unique branches used in each sample, how many of those branches are also used in at least one other sample, and how many are sample-unique; a companion file reports, for every (hub, target) pair, the branch fraction and target-level expression in each sample. Comparing entropy values across samples reveals condition-specific shifts in splicing complexity — a decrease in entropy suggests consolidation toward fewer dominant isoforms, while an increase indicates diversification of splice-site usage.

### Read clustering, splice-site indexing, and transcript classification

The `query` and `discover` subcommands share a coarse-to-fine matching routine. Input records — individual transcripts for `query`, aligned reads for `discover` — are reduced to their ordered splice-junction signatures. For reads, junctions are extracted from CIGAR `N` (reference-skip) operations and the primary-alignment filter is applied at `bam_- reader` level. Reads are then clustered by a strand-aware sort-and-sweep over their (strand, junction count, first donor) key; any two reads whose junction lists agree within `--fuzzy-tolerance` bp are collapsed into the same cluster.

For each cluster or input transcript, the matcher performs a spatial `intersect` against the grove to obtain candidate segments, walks each candidate via its SEGMENT-TO-EXON edge to the first exon, and then traces the EXON-TO-EXON chain, comparing query junctions against exon boundaries as it goes. To classify individual donor and acceptor positions as known or novel, the matcher first walks the entire grove once to hash every exon’s 5^*′*^ and 3^*′*^ end into two global sets `(known_donor_sites and known_acceptor_sites);` this indexing step operates directly on a pre-built `.ggx` index and does not require any build-time caches.

Query transcripts are assigned to SQANTI-like structural categories following Pardo-Palacios et al. (2024): Full Splice Match (FSM) when every junction matches a candidate segment within tolerance, Incomplete Splice Match (ISM) when the query is a contiguous subsequence of a segment, Novel In Catalog (NIC) when all donors and acceptors are known but their combination is not, Novel Not in Catalog (NNC) when at least one donor or acceptor is not in the splice-site lookup, and the GENIC-INTRON, GENIC-GENOMIC, ANTISENSE, INTERGENIC, and AMBIGUOUS categories for progressively weaker forms of evidence. ISM subcategories (*prefix, suffix, internal*), NIC subcategories (*combination, intron retention, alternative splice*), and NNC subcategories (*novel donor, novel acceptor, novel both*) are resolved from the same traversal. Discovery clusters with no compatible segment but sufficient read support are inserted as new segments tagged with `atroplex` as their source.

### Differential transcript usage

Given a two-group contrast specified on any `sample_info` field (e.g., `--contrast treated:control`), the `query` subcommand computes, for every gene, a *G* ×*S* count matrix where *G* is the number of segments of the gene and *S* = 2 is the number of groups. Within each group, the segment counts are obtained by summing per-sample expression (or presence, if no expression is available) over all samples assigned to that group. A gene-level Pearson *χ*^2^ test on the resulting contingency table then tests whether the segment usage proportions differ between groups; *p*-values are corrected across genes by the Benjamini–Hochberg procedure at a user-supplied FDR level (default 5%). Genes with fewer than two segments are excluded from testing.

---

**Figure S1.**
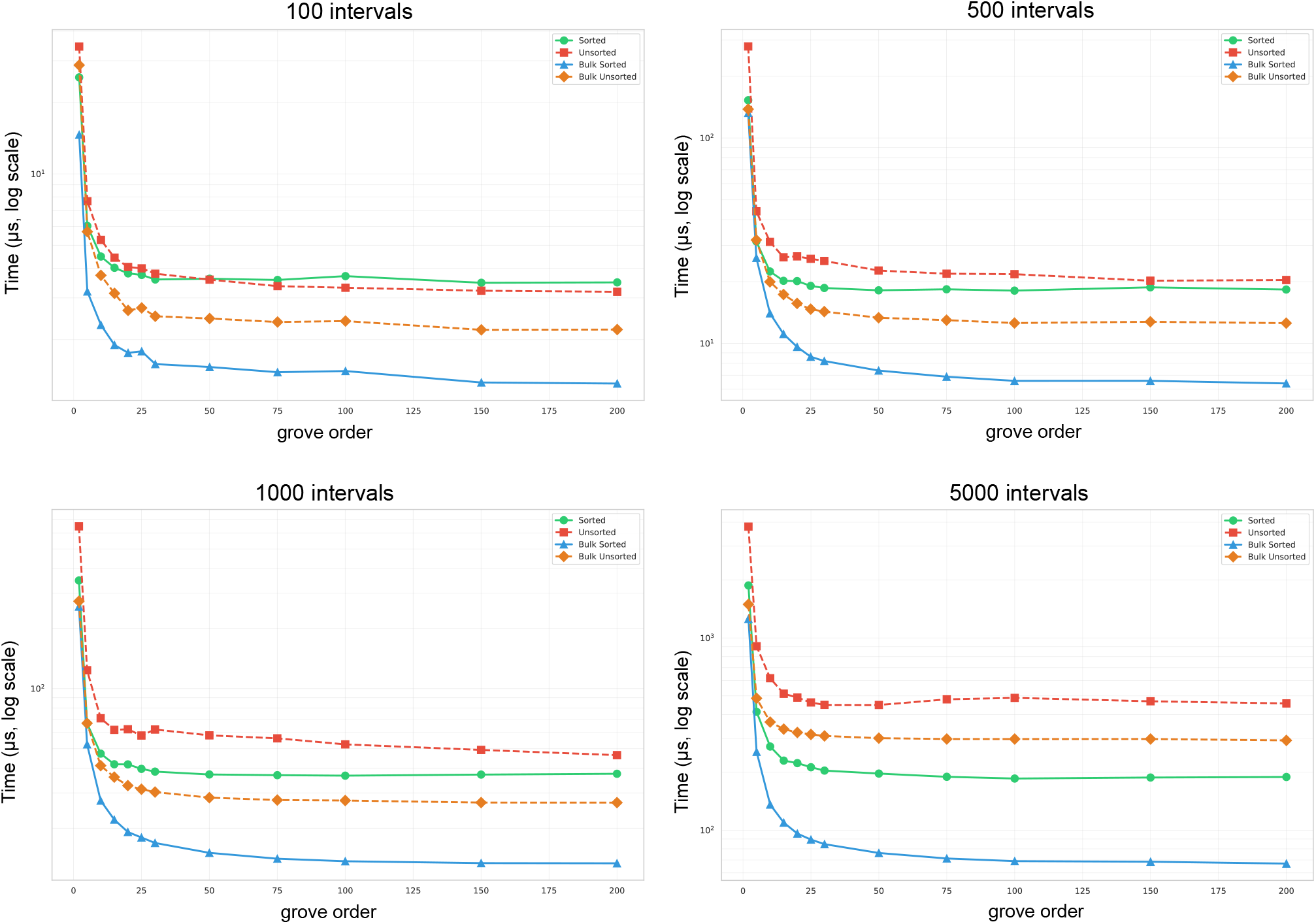
Benchmarking genogrove index construction.

**Figure S2.**
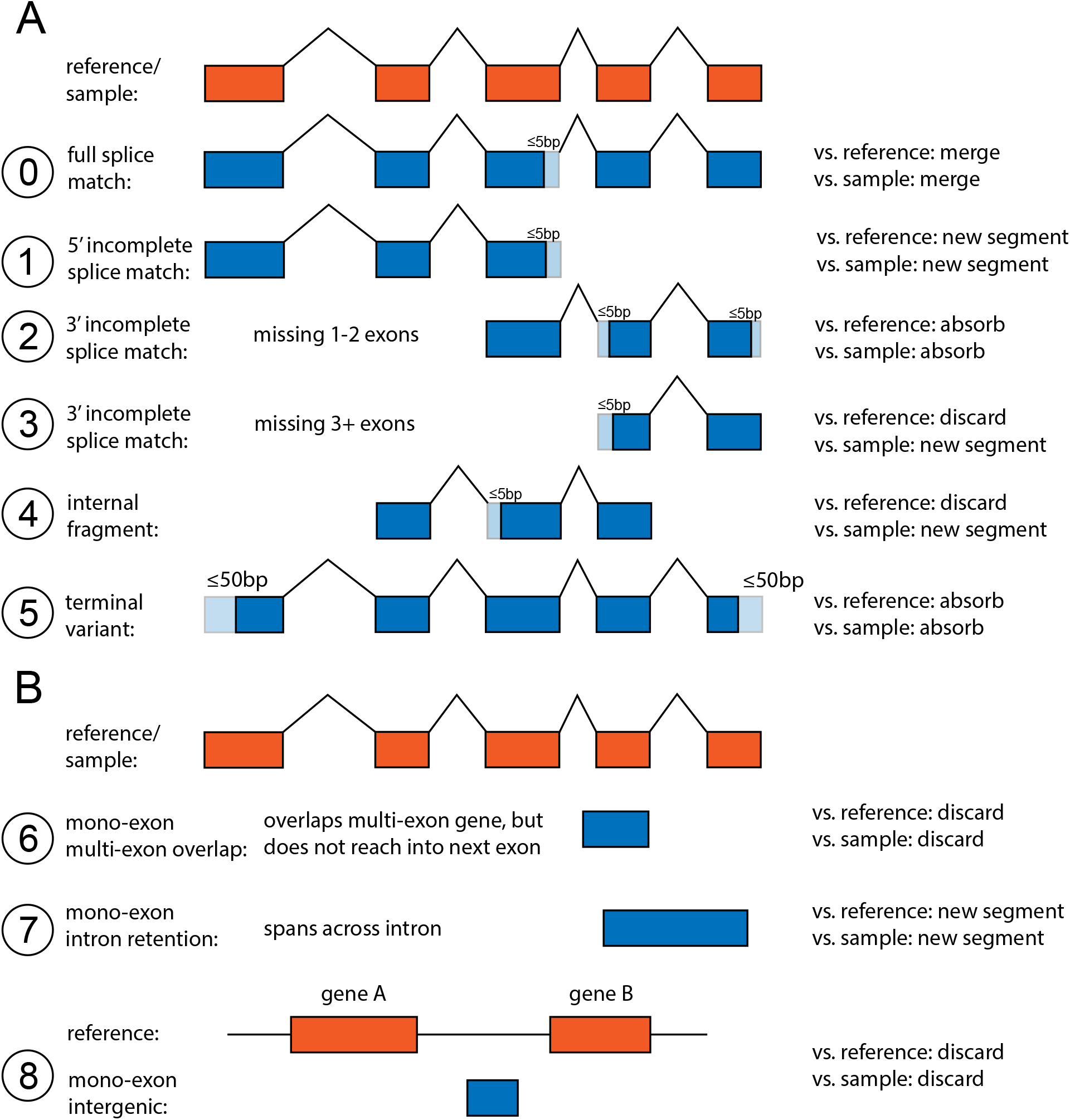
Absorption rules for multi-exon transcripts during index creation. (**A**) Transcripts are processed within each gene locus. Rule 0 merges exact structural matches (with 5 bp fuzzy tolerance). Rules 1–4 classify contiguous exon-chain subsets: 5^*′*^ ISMs (Rule 1) are retained as potential alternative start sites; 3^*′*^ ISMs missing 1–2 exons (Rule 2) are absorbed as likely RT dropout artefacts; 3^*′*^truncations missing 3+ exons (Rule 3) and internal fragments (Rule 4) are dropped when matching a reference annotation but retained when matching another sample assembly. Rule 5 absorbs terminal variants sharing the same intron chain but differing by less than 50 bp at transcript boundaries. Annotations are always processed before samples to establish reference segments as absorption targets. (**B**) Absorption rules for mono-exon transcripts. Rule 6 drops mono-exon transcripts that overlap a multi-exon gene without spanning an intron boundary, as these are likely fragmented or degraded reads. Rule 7 retains mono-exon transcripts that span from one exon into the next across an intron, representing potential intron retention events. Rule 8 drops mono-exon transcripts with no gene overlap, filtering intergenic noise. All three rules apply identically regardless of whether the parent is a reference annotation or sample assembly.

**Figure S3.**
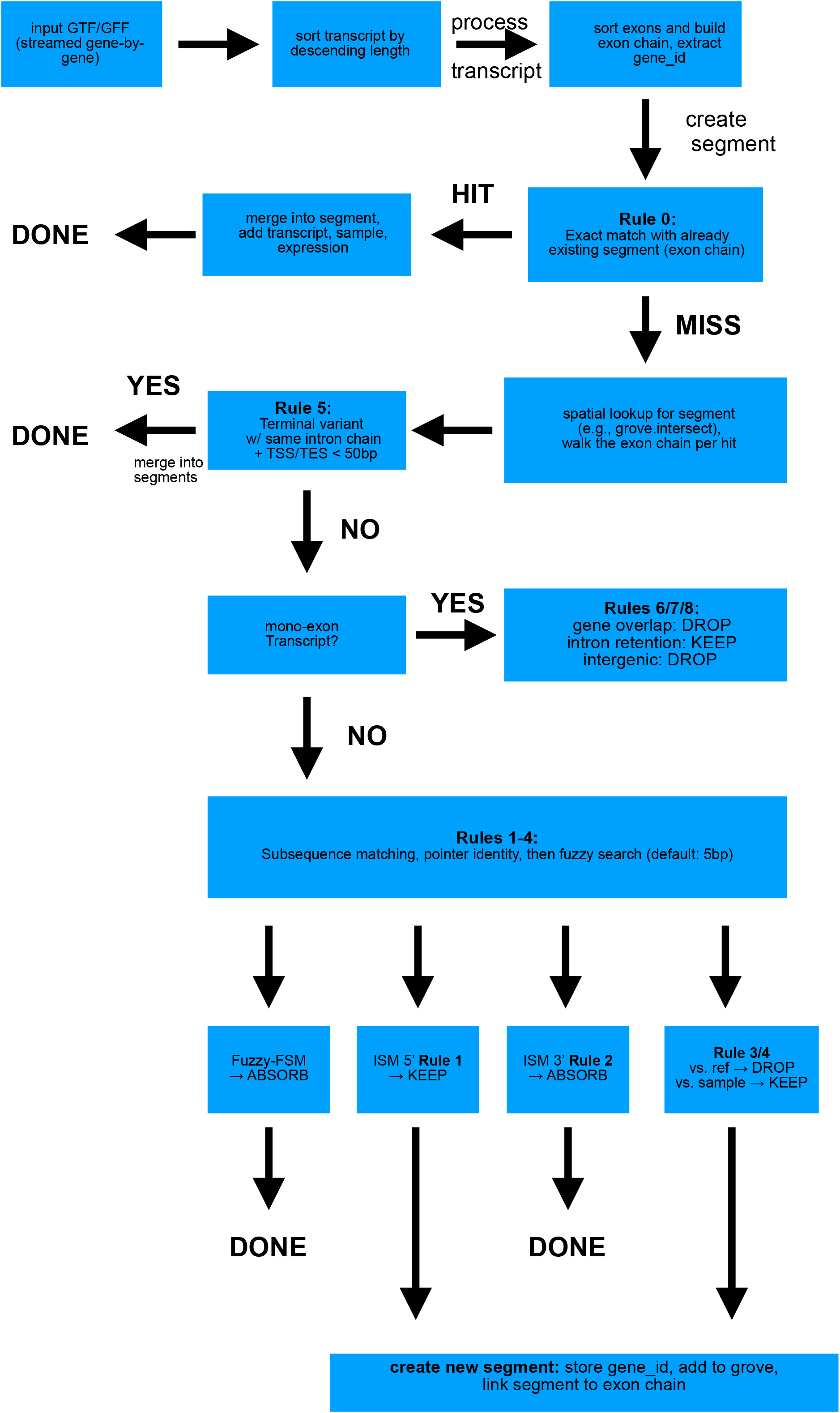
Decision flow for absorption and segment creation. Each incoming transcript is first probed for an exact structural match against the existing index; on a hit, the matching segment′s metadata is updated in place. On a miss, the spatial index returns candidate segments overlapping the transcript on the same strand, which are then evaluated against the absorption rules of Supplementary Figure S2. Transcripts that pass all checks without matching produce a new segment, after which a reverse-absorption pass merges any previously inserted segments that are contiguous subsequences of the new one.

**Figure S4.**
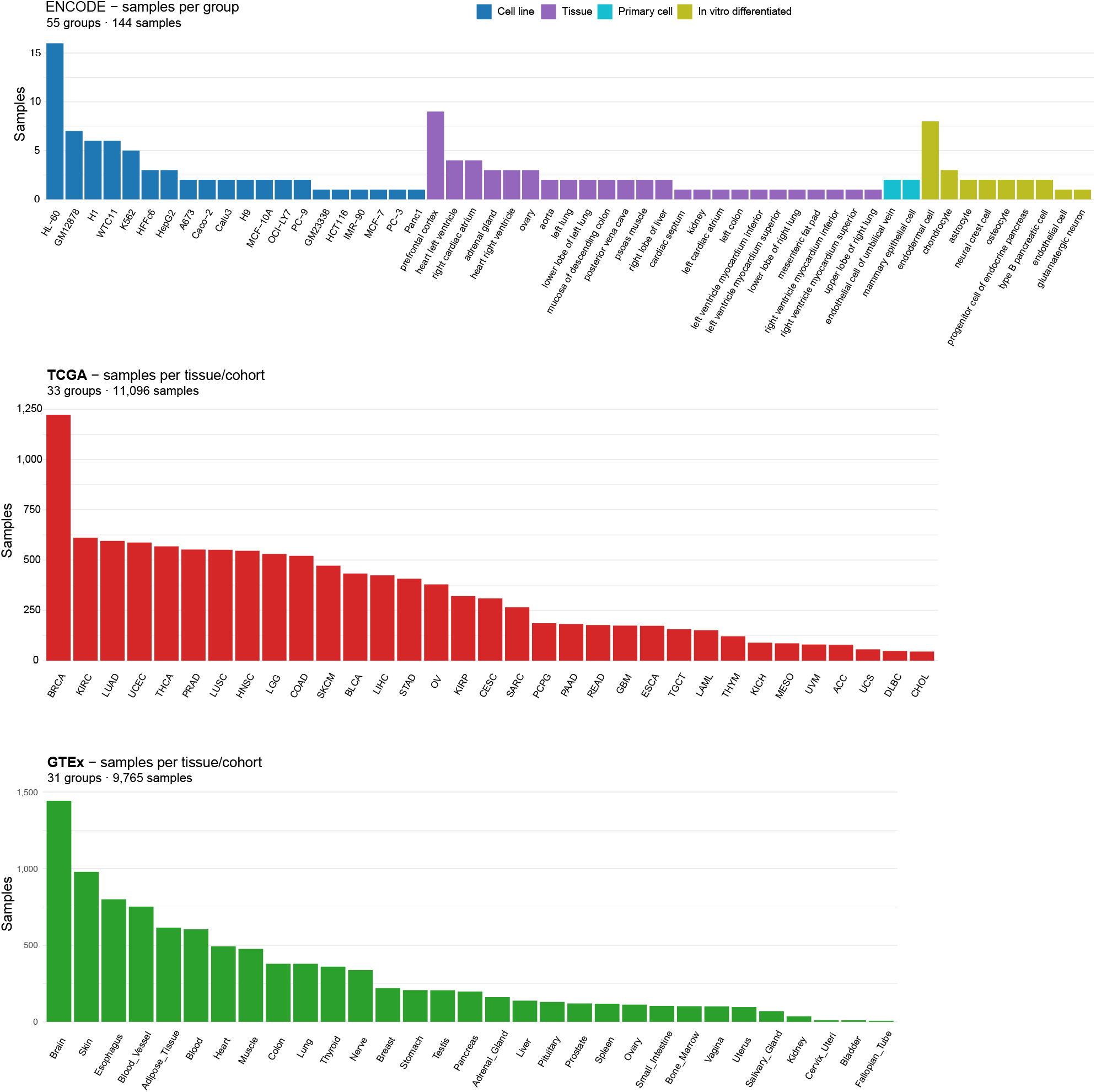
Composition of the 21,005-sample pan-transcriptome cohort. Samples are stratified by source (ENCODE4 long-read, GTEx, TCGA), tissue or cancer type, and assembly pipeline. Reference annotations (GENCODE v49, ENCODE4_LR) are processed before experimental samples during index construction.

## Notes

https://github.com/ylab-hi/atroplex

### Competing Interest Statement

The authors have declared no competing interest.

